# Optimizing automated classification for zooplankton in coastal conditions: the impact of model selection, imaging instruments, and colour information

**DOI:** 10.64898/2026.07.09.733739

**Authors:** Pieter D. L. Hovenkamp, Lodewijk van Walraven, Anouk Ollevier, Dick van Oevelen, A. Frank van der Stappen

## Abstract

The advancement in deep learning techniques has made Convolutional Neural Networks (CNNs) a powerful tool for the fully automated classification of zooplankton images. In this study, we systematically investigate how network selection, colour information and differences in imaging instruments affect the classification of zooplankton images by comparing multiple state-of-the-art CNNs on images of zooplankton and marine snow from the in situ Continuous Particle Imaging and Classification Sensor (CPICS), Video Plankton Recorder (VPR), In Situ Ichtyoplankton Imaging System (ISIIS), and the on-board Plankton Imager (Pi-10). With differences between models of 7.8 to 19% in F1-score, we find that model selection strongly affects the classification performance, with EfficientNetV2S showing the most reliable overall performance. Moreover, differences between model architectures are largest for the least abundant classes (*<*100 labeled images), which implies that when these are present, careful model selection is most beneficial. The high image quality of the Pi-10 strongly increases the performance for the least abundant classes compared to the other instruments. In addition, we find a significant correlation (*r* = 0.597) between ImageNet the performance and F1-score on zooplankton images, which implies that more generally, a model that performs well on ImageNet will perform well for zooplankton classification. Colour information increases the F1-score of the best performing classifier with 2.8%, but provides a stronger benefit (25% F1-score) for classes with *<*100 images. The overall performance increase of colour information is less than expected and questions the advantage of recording colour information for zooplankton.

## Introduction

The strong response of zooplankton to local and global environmental changes (Richardson, 2008; Mackas and Beaugrand, 2010; Beaugrand et al., 2013) that are occurring this century has led to increased interest in plankton monitoring. This interest has driven the development of various plankton imaging instruments (see Lombard et al. (2019) for an overview) that enable us to collect large amounts of data on zooplankton distribution at a finer spatial and temporal scale than with traditional net sampling (Irisson et al., 2022). Meanwhile, classifying these vast amounts of data is costly and requires taxonomic expertise that is declining (Cotterill and Foissner, 2010), and therefore fully automated image classification of plankton images is much desired. Advances in deep learning methods have made Convolutional Neural Networks (CNNs; LeCun et al., 2015; Krizhevsky et al., 2017) an accessible and powerful tool within the field of zooplankton imaging. This relatively new technique, which is a form of so-called ‘deep learning’, was quickly adopted for the application of image recognition of plankton images, first by Orenstein et al. (2015) on phytoplankton data. This was soon followed by many authors (Al-Barazanchi et al., 2018; Luo et al., 2018; Ellen et al., 2019; Guo et al., 2021; Bi et al., 2024; Yuan et al., 2024; Slocum and Penta, 2025) who, among others, have successfully applied CNNs to fully automated classification of in situ images of zooplankton.

With the growing number of imaging instruments that are designed for specific purposes and with the ongoing evolution of the quality of images that are produced, there is a high variability in plankton images among different instruments. Some notable aspects are the size range of the targeted organisms, the illumination technique or whether the instrument produces colour or grayscale images. For example, the Continuous Particle Imaging and Classification Sensor (CPICS; Gallager, 2016) and the Video Plankton Recorder (VPR; Davis et al., 2005) collect dark field images versus white field images collected by the In Situ Ichthyoplankton Imaging System Deep-focus Particle Imager (ISIIS-DPI; Cowen and Guigand, 2008), Plankton Imager (Pi-10; Pitois et al., 2018) and the Underwater Vision Profiler (UVP; Picheral et al., 2010). This makes it difficult to interchange data sets or classifiers as these are tailored to specific instruments (Orenstein and Beijbom, 2017; Luo et al., 2018; Batrakhanov et al., 2024). In addition, variability in species distribution among regions and seasons can severely reduce the performance of a previously trained classifier (Moreno-Torres et al., 2012; Gonźalez et al., 2017). Consequently, as new instruments are developed or introduced in new regions and the number of annotated zooplankton images grows, reusing previously trained networks is not beneficial. With automated image analysis increasingly being applied, training a CNN is therefore becoming a standard task in the field.

Since AlexNet, a pioneering CNN model, outperformed other types of classifiers in the ImageNet Large Scale Visual Recognition Challenge of 2012 (Krizhevsky et al., 2017), much research effort has been put into further improving the performance of CNN architectures. This was done for example by creating heavier networks with more parameters, by introducing new network structures such as inception (Szegedy et al., 2014) or residual modules (He et al., 2016), by optimising performance for real-time classification (MobileNet; Sandler et al., 2018) or by optimising - given computational limitations - the relation between input size, network width and number of layers (EfficientNet; Tan and Le, 2021). This has led to a large, and growing, number of models that are available ‘out-of-the-box’ (e.g. in platforms such as Keras or Hugging Face) which can be used for plankton image classification. The performance of these models is benchmarked on ImageNet (Deng et al., 2009), an open-source, large-scale database of annotated images of varying object types (e.g. vehicles, animals, furniture). However, ImageNet and plankton images are markedly different and this raises the question whether model architectures that perform better on ImageNet are also more suitable for classifying plankton images.

Lumini and Nanni (2019) compared the performance of various networks on grayscale plankton images from WHOI-Plankton (Orenstein et al., 2015), ZooScan (Gorsky et al., 2010) and ISIIS-DPI in order to determine the best performing model or ensemble of models. Since then, new state-of-the-art models have emerged and additionally, the presence of colour information might improve the performance of some architectures more than others. Kyathanahally et al. (2021) compared the performance of various modern CNNs on lake zooplankton with the Scripps Plankton Camera (Orenstein et al., 2020), and achieved classification accuracy of 97.9% by ensembling multiple networks. However, ensembling many CNNs vastly increases model inference time and with growing amount of images that can be collected per deployment (e.g. Schmid et al., 2023b; Pitois et al., 2025, Hovenkamp et al., in press), computational time poses a limit to this possibility. In addition, an apparent trend is that better individual models also lead to better model ensembles. Although new techniques, such as (ensembles of) Vision Transformers show promising results for zooplankton classification (Kyathanahally et al., 2022; Maracani et al., 2023; Nanni et al., 2023; Yue et al., 2023; Li et al., 2025), these are generally computationally heavy (Rubbens et al., 2023; Nanni et al., 2025) and the widely available CNNs are still a common choice in ecological studies. Therefore, the question which individual CNN models perform best for zooplankton classification remains relevant. Moreover, we extend the previous studies on this topic by linking performance differences between CNN models to the instrument type and characteristics of training classes.

In addition, with the technical advancement of imaging hardware, several instruments have been developed that acquire colour images. This increases the amount of generated image data compared to grayscale images, and since the amount of data that can be stored per unit of deployment time often is limited, any increase in generated data per image inherently compromises the maximum frame rate, and thus sampling volume, of an instrument. A growing number of instruments use a silhouette illumination technique, such as the ISIIS-DPI and Pi-10. This technique is not able to record colour information accurately: ISIIS-DPI images are stored as grayscale and those of Pi-10 as RGB but with barely visible colour. However, it does enable a sampling volume that is substantially higher than a colour imaging technique (Cowen and Guigand, 2008). Given the negative relationship between organism size and abundance (Sheldon et al., 1972), it is the instrument’s sampling volume that determines the upper size limit of planktonic organisms that can be accurately sampled (Lombard et al., 2019). Thus, the presented trade-off between recording colour and sampling volume raises the question whether colour information is beneficial for the automated classification of plankton with CNNs.

In this study, we address and connect the issues of CNN model selection and the relevance of storing colour information for zooplankton image classification. We take a systematic approach and compare various well-established techniques that are used in field studies in order to identify which CNN classifier gives the best overall performance, understand the variability in classifier performance across different classes and instruments, and investigate whether incorporating color information enhances CNN performance compared to using grayscale images. In addition, we explore the influence of network choice, colour information, and instrument differences on the relationship between the size of training classes and classification performance.

## Methods

### Imaging instruments and data sets

We utilised training data from four different zooplankton imaging instruments: the Continuous Particle Imaging and Classification Sensor (CPICS, 10,275 images), the Video Plankton Recorder (VPR, 37,239 images), the In Situ Ichtyoplankton Imaging System (ISIIS-DPI, 13,417 images), and the Plankton Imager (Pi-10, 16,684 images).

### Continuous Particle Imaging and Classification Sensor (CPICS)

The CPICS-1000-e (Gallager, 2016) is an in situ open-flow imaging device that acquires darkfield colour images of planktonic organisms and particles (Figure 1a - f). With a 0.9× magnification lens and an image resolution of 2,736 × 2,192 pixels, the field of view is 15 × 11 mm and the pixel resolution is 4.54 µm, and in this study, the depth of field was factory-calibrated as 2mm. The CPICS is equipped with an on-board segmentation procedure that detects and stores image segments, so-called Regions of Interest (ROIs), of individual particles or organisms, irrespective of the ROI’s content.

**Figure 1:**
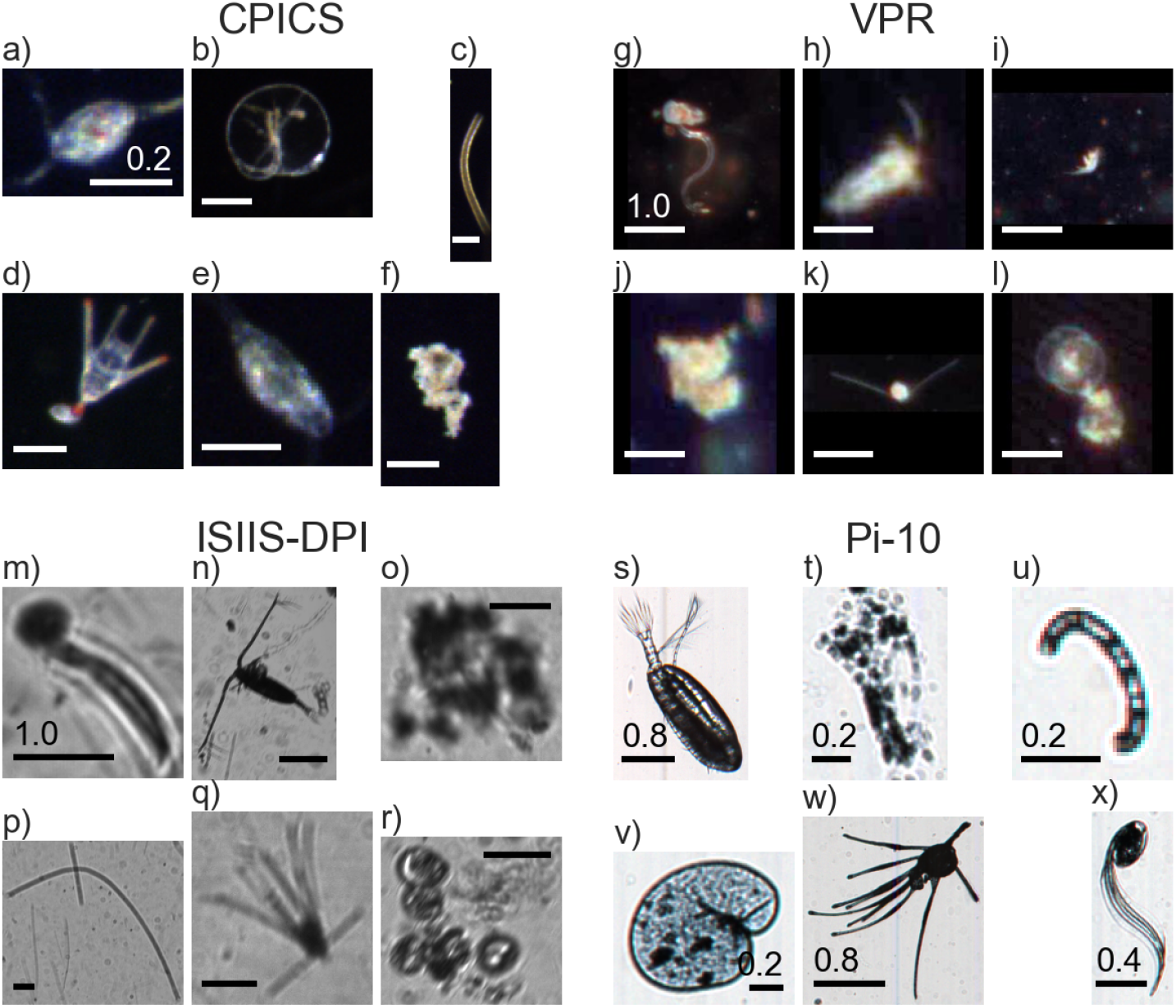
Example Regions of Interest (ROIs) from the Continuous Particle Imaging and Classification Sensor (CPICS, a - f), Video Plankton Recorder (VPR, g - l), In Situ Ichtyoplankton Imaging System (ISIIS-DPI, m - r) and Plankton Imager (Pi-10, s - x) for some example classes in our learning sets. Scale bars denote ROI size in mm and are of equal true size per instrument, except for the Pi-10 where the size is indicated per ROI. The corresponding class names are: **a)** Copepod Cyclopoid; **a, l, r, v)** *Noctiluca scintillans*; **c, p)** Diatom chain; **d, k, q, w)** Echinoderm pluteus; **e, h)** Copepod Calanoid; **g, m)** Appendicularia; **x)** Appendicularia *Oikopleura spp.*; **i)** Copepod Harpactacoid; **f, j, o, t)** Detritus; **n)** Copepod Calanoid (feeding); **s)** Copepod; **u)** Diatom chain (loop).

Data with the CPICS were collected in Lake Grevelingen in the southwest of the Netherlands. Lake Grevelingen is a former estuary that was dammed off in 1971 and still being a salt-water system, is only connected to the neighbouring North Sea via a narrow sluice. 1,805,477 ROIs were collected at 11 sites during a monthly sampling campaign from March 2021 to November 2021 and in March 2022.

Of the collected ROIs, a considerable amount were too unsharp for a taxonomist to identify as plankton. Therefore, Canny’s edge detection algorithm (Canny, 1986) was used (using *σ* = 3, and as low and high threshold respectively 10% and 20% of the maximum pixel intensity) to remove all ROIs that contained no detected edge pixels. The remaining ROIs were labeled into either one of 53 taxonomic plankton classes or one of the descriptive classes ‘non-living’, ‘bubble’, ‘out of focus’, ‘unknown sphere’ or ‘unknown living’. Only classes with a minimum number of 20 ROIs were used, and classes’out of focus’ and’unknown living’ ROIs were excluded due to heterogeneity of these classes and the lack of an ecological purpose. This results in a labeled set of 10,275 ROIs among 29 classes, of which 26 classes (7,435) of living plankton (Figure 2a).

**Figure 2:**
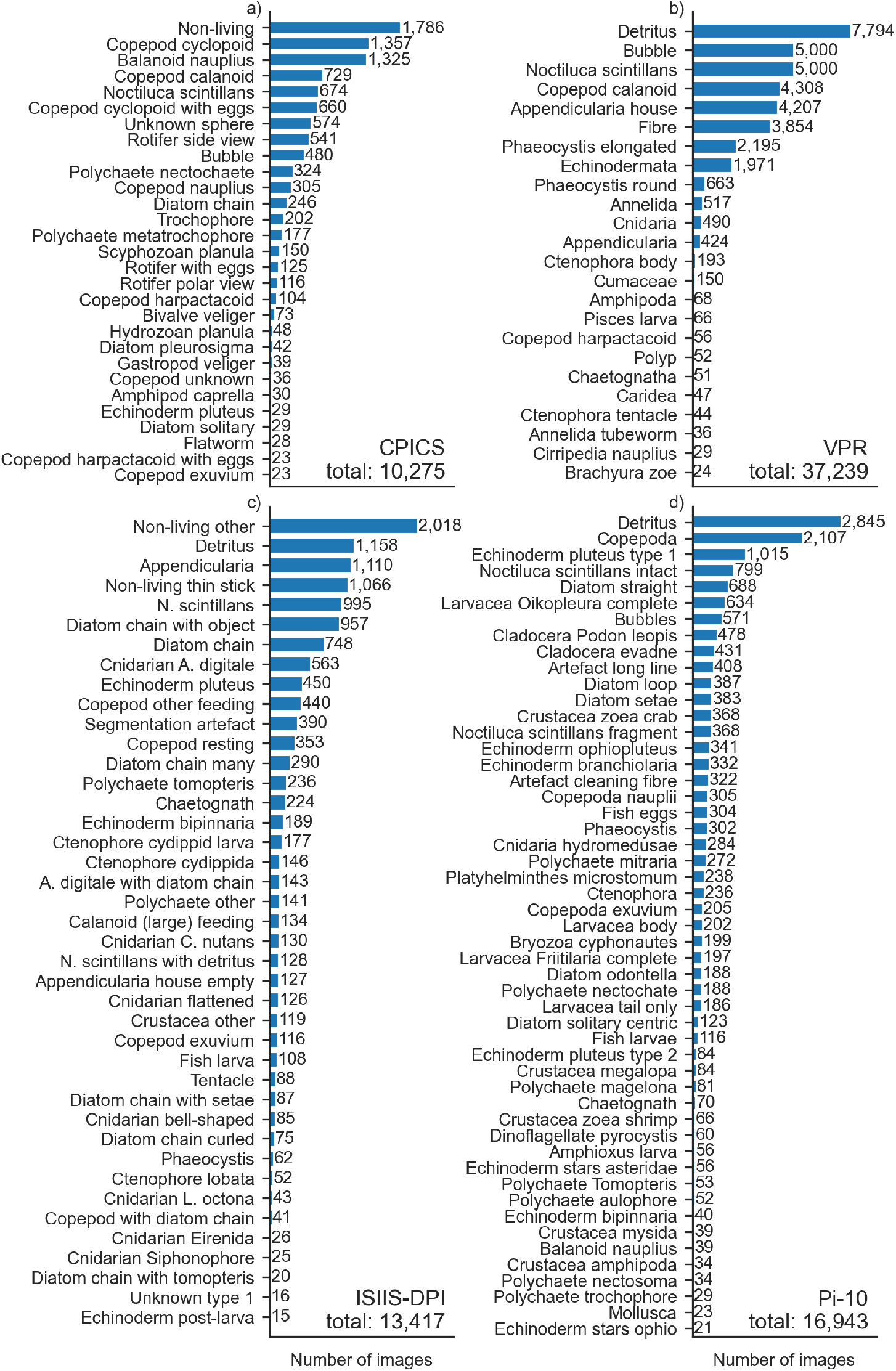
Number of ROIs in the learning sets of CPICS (a), VPR (b), ISIIS-DPI (c) and Pi-10 (d).

### Video Plankton Recorder (VPR)

A Real Time VPR acquires in situ darkfield colour images of plankton and marine snow (Figure 1g - l) with a (full-frame) image resolution of 1,380 × 1,034 and four possible magnification settings that determine the field of view, being 8.8 × 6.6 mm, 20.8 × 15.2 mm (the majority in this study), 33.8 × 25.5 mm or 46.5 × 34.5 mm. Like the CPICS, ROIs from the full-frame images are detected and stored in real-time.

Data with the VPR were collected at various sites in the Belgian North Sea between 2019 and 2021, from which we used a publicly available data set of 175,000 labeled ROIs into 32 classes (Flanders Marine Institute (VLIZ) Belgium, 2023, https://doi.org/10.14284/607). Only classes with a minimum number of 20 ROIs were used, and from the abundant classes ‘detritus’ (115,249), ‘bubbles’ (8,063) and ‘*Noctiluca scintillans*’ (24,594) a random sample of 5,000 ROIs was used. Furthermore, the classes ‘detritus’ and ‘detritus flake’ were merged, and similarly to the CPICS set, the classes ‘blurry’, ‘rest’ and ‘unknown’ were excluded. This resulted in a data set of 37,239 ROIs into 24 classes, of which 21 classes of living plankton (20,591 ROIs, Figure 2b).

### In Situ Ichthyoplankton Imaging System Deep-focus Particle Imager (ISIIS-DPI)

The ISIIS-DPI (Cowen 2008) is a shadowgraph imaging system of which data was used from an area-scan configuration with a pixel resolution of 17 µm and a (full-frame) image area of 8.83 cm (5,328 pixels) × 7.64 cm (4,608 pixels) (Figure 1m - r). The system stores full-frame images, and thus a post-processing segmentation procedure is required to extract the possibly hundreds of ROIs that can be present on a single full-frame image. The ISIIS-DPI data we used in this study were collected in the UK North Sea in June 2023, and data collection, segmentation, and manual labeling of ROIs was described in Hovenkamp et al. (in press). As in this study, our ISIIS-DPI set contained 13,417 labeled ROIs into 41 classes with a minimum number of 15 ROIs per class, of which 37 classes (9,175 ROIs) were living (Figure 2c).

### Plankton Imager (Pi-10)

The Pi-10 is an on-board instrument that continuously images surface water that is pumped through a flow cell (Pitois et al., 2018). With a line scan camera, it acquires white-field images (Figure 1s - x) in which ROIs are detected in real-time and a maximum of (in this study) 10,000 ROIs per minute are stored. The Pi-10 data that we used in this study were collected in June 2023 in the Dutch North Sea, during which object detection thresholds were such that only ROIs sized from 200µm to 2cm were stored. Data collection and labeling is further described in Van Walraven et al. (2023, in Dutch). From this data set, 16,943 ROIs into 51 classes were used with a minimum of 20 ROIs per class, of which 12,797 ROIs among 47 classes were living plankton (Figure 2d).

### Training data

All manually annotated sets, or learning sets, were divided into 70% training, 15% validation and 15% test data via a stratified split, which means that an equal proportion per label was used. Following common practice in supervised machine learning, a model is trained on the training set, and the validation set serves as an independent performance assessment throughout the training procedure in order to prevent overfitting. Decisions during the training procedure, such as selecting the best training epoch, are taken based on the validation set. As this can still lead to some degree of overfitting, the test set is used for the final performance assessment and is completely independent of the training procedure.

Note that there are intrinsic differences among the available data sets in i) the number of ROIs (hereafter referred to as ‘image’), ii) number of classes and iii) the distribution of images among classes. This situation is representative for real-life applications. For example, recent zooplankton field studies with automated image analysis (e.g. Schmid et al., 2020; Bi et al., 2022, 2024; Panäıotis et al., 2023; Yuan et al., 2024; Gastauer and Ohman, 2025) used 12 (Yuan et al., 2024) to 58 (Panäıotis et al., 2023) classes with the number of ROIs per class varying e.g. from 3 to 20,212 (Bi et al., 2024) and 46 to 608 (Yuan et al., 2024). In addition, there is intrinsic variation in the taxonomic resolution between the classes of the different annotated sets: for example, the Pi-10 contains 7 classes of echinodermata versus 1 for the VPR. These differences in taxonomic resolution are attributed to differences in image quality (as discussed later) and do not necessarily imply that the inter-class similarity between the learning sets is different.

### Experimental set-up

We conducted three experiments in which we trained CNN architectures on the data sets from the various instruments:

- in experiment 1, we trained 18 CNN architectures on the CPICS set in order to determine the architecture with the best performance on this data set; For this, 18 modern CNN classification architectures (Table A2) from 9 model families were selected that were pre-trained on ImageNet and that are widely used in the literature with an ImageNet top-1 accuracy of 71.3 83.9%.
- in experiment 2, the best models from experiment 1 were trained on the learning sets from the VPR, Pi-10 and ISIIS-DPI to extend the performance comparison of experiment 1 to the other instruments. In addition, we used the results to assess performance differences between model architectures and instruments;
- in experiment 3 we created an additional learning set, called CPICS_gray_, by converting the images from the CPICS learning set to grayscale (see Figure A1 for example images), and we trained the best models from experiment 1 on this grayscale set. In this set, we kept the division between training, validation and test data equal to its RGB equivalent (referred to as CPICS_RGB_). By comparing the classifiers trained on CPICS_gray_ to those trained on CPICS_RGB_, we assessed whether colour information improves the classification accuracy of zooplankton images.

Experiment 1 was carried out on the CPICS set only, because the extensive training procedure with hyperparameter tuning was too computationally costly to repeat for all instruments. Therefore, based on the results of this experiment, we selected the best performing models for the consecutive experiments 2 and 3: from five different model families we selected the models with the best performance, and regardless of performance, ResNet50V2 was added to this model selection, as ResNet50(V2) is commonly used in the zooplankton literature (e.g. Guo et al., 2021; MacNeil et al., 2021; Schanz et al., 2023; Pitois et al., 2025).

### Training procedure

Training of all CNNs was carried out in a standardised way that includes transfer learning (Ponti et al., 2017; Rubbens et al., 2023) and tuning of so-called hyperparameters, the term for parameters that specify the training procedure.

With transfer learning, all trainable parameters before the final classification layers of a model are copied from an instance of the same model that was trained on a different data set, which in our case was ImageNet. The classification layers are then trained while keeping the weights of the base model, i.e. all layers before the classification layers, frozen. During a subsequent fine-tuning phase, the model is further trained with the weights of the complete model being trainable. Transfer learning is commonly used for zooplankton classification, because it leads to better performance with fewer training images for both zooplankton (Orenstein and Beijbom, 2017; Maracani et al., 2023; Ellen and Ohman, 2024) and non-zooplankton applications (Kornblith et al., 2019). In our study, initial trials showed that transfer learning on our data led to better model performance and faster training, and that performance was less dependent on hyperparameters than when training randomly initialised models from scratch.

A brief summary of the training procedure that we applied in all experiments is as follows. First, hyperparameter tuning using Bayesian optimisation was done, since initial training results were observed to vary strongly between different values of learning rate and optimiser’s weight decay. The importance of hyperparameter optimisation instead of picking standard values was also shown for non-plankton images (e.g. Brigato et al., 2022). During hyperparameter tuning, we trained the base model in 7 training rounds of 30 epochs, each round with a different combination for the learning rate and the optimiser’s weight decay, and the combination of learning rate and weight decay that resulted in the highest validation accuracy was used to redo the training for up to 150 training epochs. Other hyperparameters were fixed for all training rounds at a value that is common in the literature.

In the subsequent fine-tuning phase with all model weights unfrozen, another hyperparameter search was done on the learning rate and the optimiser’s weight decay in again 7 training rounds of 30 epochs, since fine-tuning generally requires a different, lower learning rate than initial training. Finally, the best performing model from the fine-tuning round is taken as the result of the experiment. During the training and fine-tuning rounds, training was aborted as soon as the validation accuracy did not improve over a certain number of epochs to avoid overfitting. After each training phase, it was visually verified that a plateau in model performance was reached. A complete overview of all hyperparameters per training round for each experiment is given in Table A3.

Since the input size of images for the CNNs that we examined is a fixed property, we downsized images with a dimension that was larger than the required input size of the considered architecture (Table A2). Smaller image dimensions were padded (in line with e.g. Ellen et al., 2019) with black and white pixels for dark-field and white-field images, respectively. In addition, contrast enhancement was applied to the sets of darkfield images during both training and evaluation. To adjust for the strong class-imbalance between the rare and abundant classes in our learning sets, balanced class weights were used in the loss function during training. Balanced class weights for a class *i* were calculated as *w_i_* = 1*/n_i_* ∗ *N*, with *n_i_* the number of images of a class and *N* the mean number of images per class over all classes. This ensures that classification errors in rare classes were penalisd more than errors in common classes. Without such a correction, a classifier would only focus on the largest classes and ignore the rare classes due to their minor contribution to the overall loss. All of our experiments were coded in Python3 (Van Rossum and Drake, 2009) using Tensorflow (Mart‘ın Abadi et al., 2015), KerasTuner (O’Malley et al., 2019), and scikit-learn (Pedregosa et al., 2011) and performed on a Nvidia Tesla-V100-PCIE-32GB GPU.

### Performance evaluation

The performance of the classifiers was assessed by comparing the predictions against the verified labels (also called ‘ground truth’) from the test set of each data set. The accuracy is defined as the number of correct predictions in the data set divided by the total number of predictions (Eq. 1). This metric is biased towards the most prevalent classes if a test set is unbalanced, a situation that often occurs in zooplankton applications (Schröder et al., 2019; Eerola et al., 2024; Masoudi et al., 2024). Therefore, we additionally report the F1-score (Eq. 4), a class-specific metric that is commonly used and depends on the precision (Eq. 2) and recall (Eq. 3).

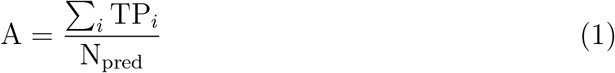

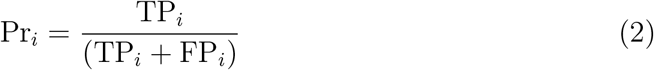

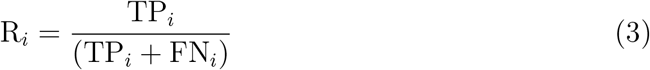

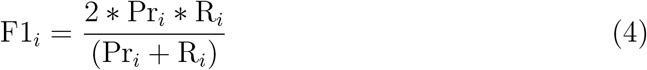

In these equations, we made use of the following additional definitions: N_pred_ is the total number of predictions, TP*_i_* is the number of true positives for a class *i* (i.e. ground truth is *i* and the prediction is *i*), FP*_i_* is the number of false positives (prediction is *i* whereas ground truth is not *i*) and FN*_i_* is the number of false negatives (ground truth is *i* whereas prediction is not *i*). The interpretation is that precision quantifies how many predictions in a class are correct (‘purity’), whereas recall measures how many of the ground truth labels of a class are actually predicted to be in that class (‘completeness’). Calculating the average F1-score over all classes results in an overall metric that equally takes into account the performance of all classes, and thereby compensates for unbalanced data sets.

## Results

### Network comparison

The training of all 18 CNN architectures on CPICS_RGB_ in experiment 1 (Figure 3) shows that the accuracy of the resulting models ranges from 68.0% (MobileNetV2) to 90.8% (EfficientNetV2S), and F1-score from 48.4% (MobileNetV2) to 79.0% (EfficientNetV2B2). For all models, the F1-score is considerably lower than the accuracy. This is to be expected: rare classes generally have a lower performance and these contribute more strongly to the F1-score than to the accuracy.

**Figure 3:**
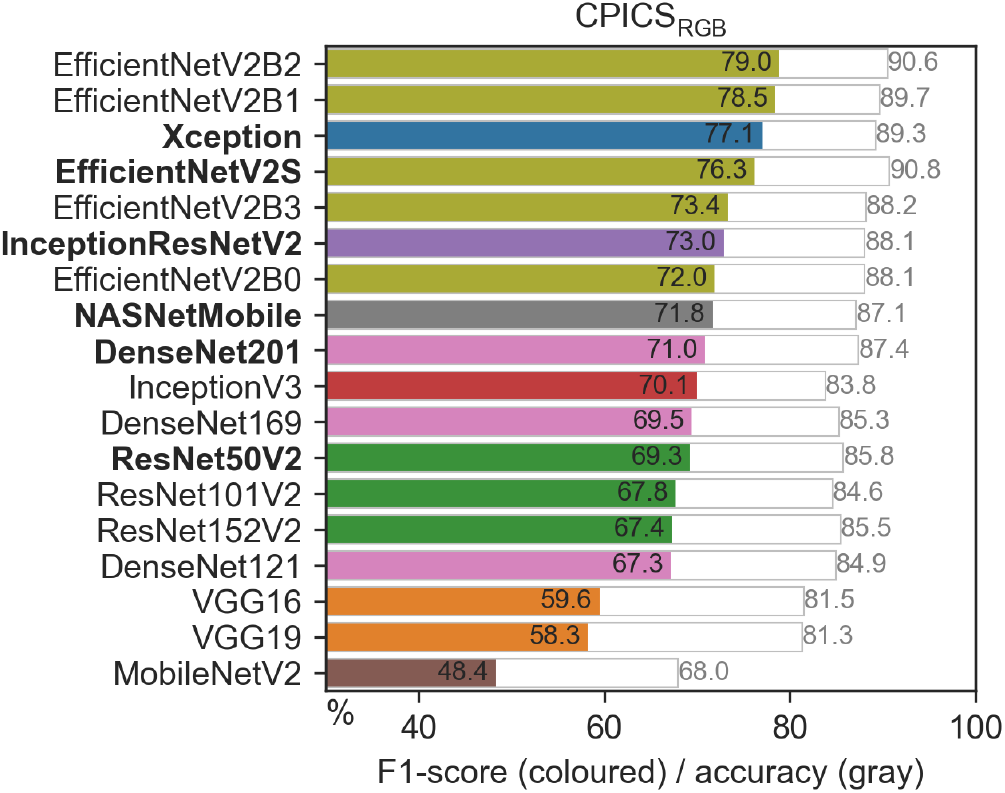
Comparison of accuracy (grey) and F1-score (coloured) of all assessed models on learning set CPICS_RGB_ (experiment 1). Models in bold were used in the subsequent experiments.

For experiment 2, we selected 5 models from different model families based on the performance on CPICS_RGB_. Evaluation in accuracy and F1-score (Figure 3) both lead to the same top-5 of model families, consisting of Xception, InceptionResNetV2, NASNetMobile, EfficientNetV2 and DenseNet. To select the best model within the EfficientNetV2 and DenseNet family, we decided to choose

EfficientNetV2S and DenseNet201, as these are the heaviest variants within the model families and thus expected to achieve the best performance. As already mentioned in the method section, we also included ResNet50V2 as a sixth model in the remaining tests, because ResNet50(V2) is commonly used in other plankton studies (Guo et al., 2021; MacNeil et al., 2021; Schanz et al., 2023; Pitois et al., 2025).

Experiment 2 show that in terms of accuracy, the performance of EfficientNetV2S (83.4 to 93.0%) is best for all considered data sets (Figure 4). Measured in F1-score, EfficientNetV2S performs best on the ISIIS-DPI (78.9%) and CPICS_gray_ (73.5%), but for the VPR, DenseNet201 performs slightly better (77.7% versus 77.5%) and for CPICS_RGB_ this is the case for Xception (77.1% versus 76.3%). For the Pi-10, NASNetMobile (89.3%) shows the best F1-score. Notably, for the Pi-10 the F1-score of ResNet50V2 (81.5%) is considerably lower than for the top-5 models (87.6 to 89.3%), but the performance difference within the top-5 models is only 1.5% (accuracy) and 1.7% (F1-score). This is smaller than for the other instruments, where the difference within the top-5 ranges from 3.3% (VPR) to 5.8% (CPICS_gray_) in accuracy and 6.1% (CPICS_RGB_) to 10.6% in F1-score (VPR). Finally, we find a significant relationship between the ImageNet top-1 accuracy from the literature and our models’ F1-score (Pearson’s *r* = 0.597 and *p* = 0.0001, Figure 5).

**Figure 4:**
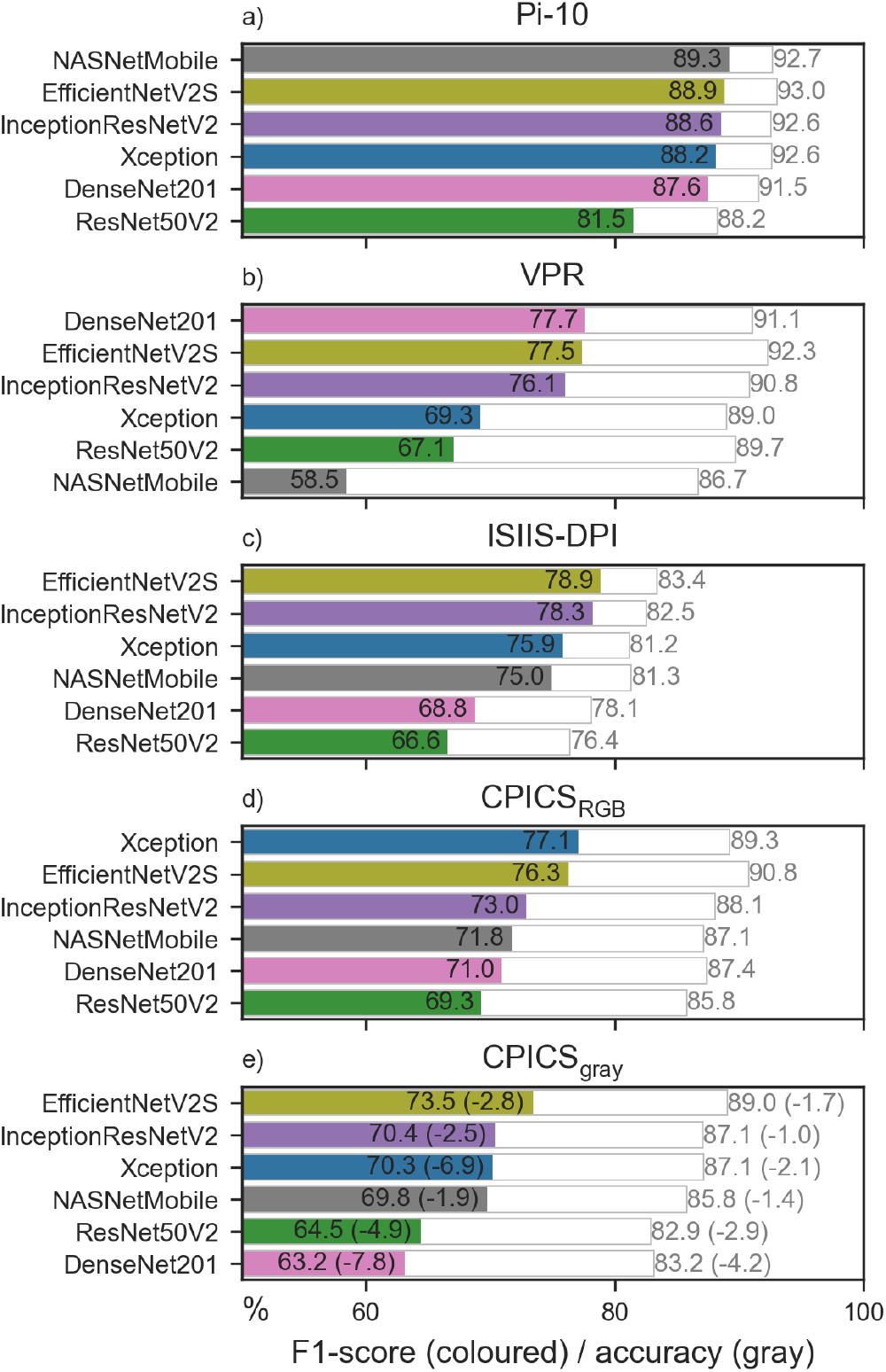
Comparison of accuracy (grey) and F1-score (coloured) of the top-6 models on learning set Pi-10 (a), VPR (b), ISIIS-DPI (c), CPICS_RGB_ (d), and CPICS_gray_ (e). The numbers in parentheses in (e) denote the difference between performance on CPICS_gray_ and CPICS_RGB_.

**Figure 5:**
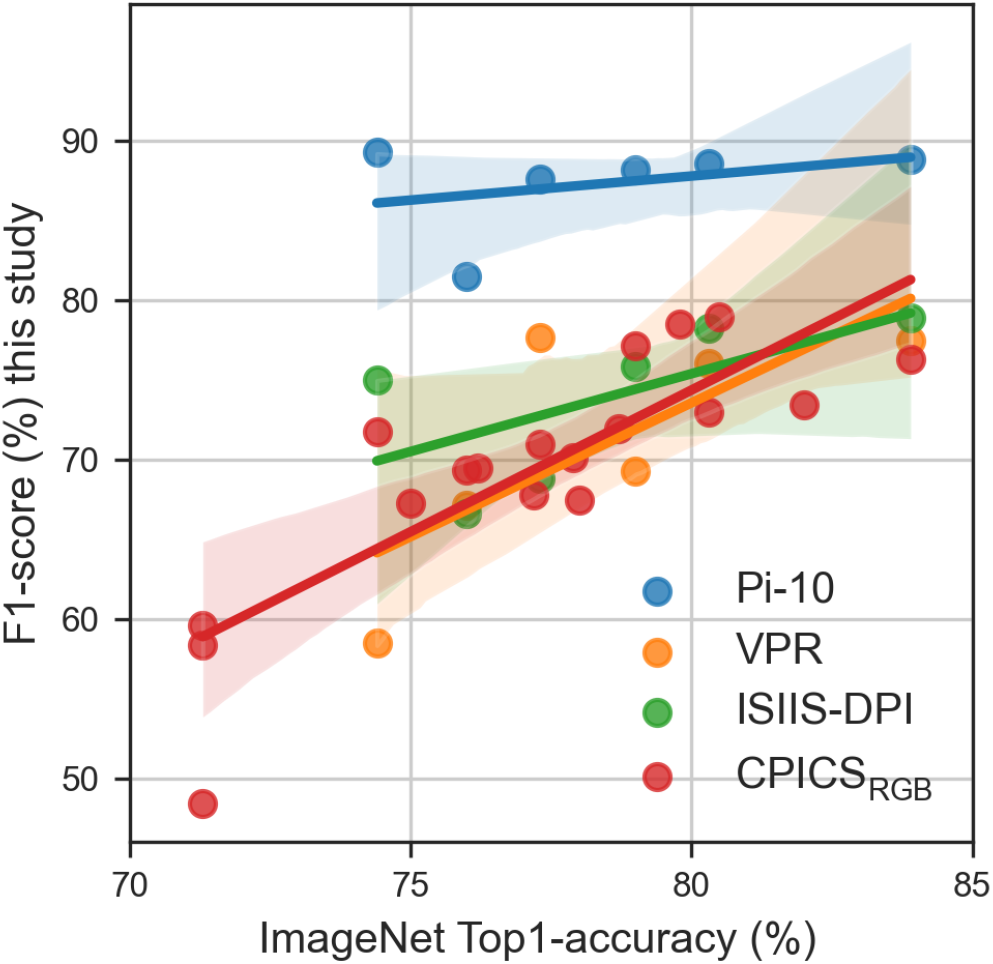
Comparison of the accuracy on ImageNet (Table A2) versus the F1-score on our learning sets for each assessed model. Note that the experiments with CPICS_RGB_ include more models (16) than with the other instruments (6).

### Class-specific performance

The model performance per taxonomic class in experiment 2 (Figure 6) shows that for all learning sets, the performance difference between models decreases with increasing class size. This is particularly the case for the VPR, ISIIS-DPI and CPICS_RGB_ sets, where for classes with less than 100 images differences in F1-score range from ∼20 to 40%, whereas for the highest number of images these differences are in the order of 5%. A notable difference between the Pi-10 set and the other instruments is that the F1-score for the training classes with few images (30 to 100) is higher (*>*∼ 80%) than for the other instruments. Also, for the Pi-10 learning set, differences between models are smaller (∼5 to 15%) than for the other instruments (∼5 to 40%) for classes with more than 30 training images.

**Figure 6:**
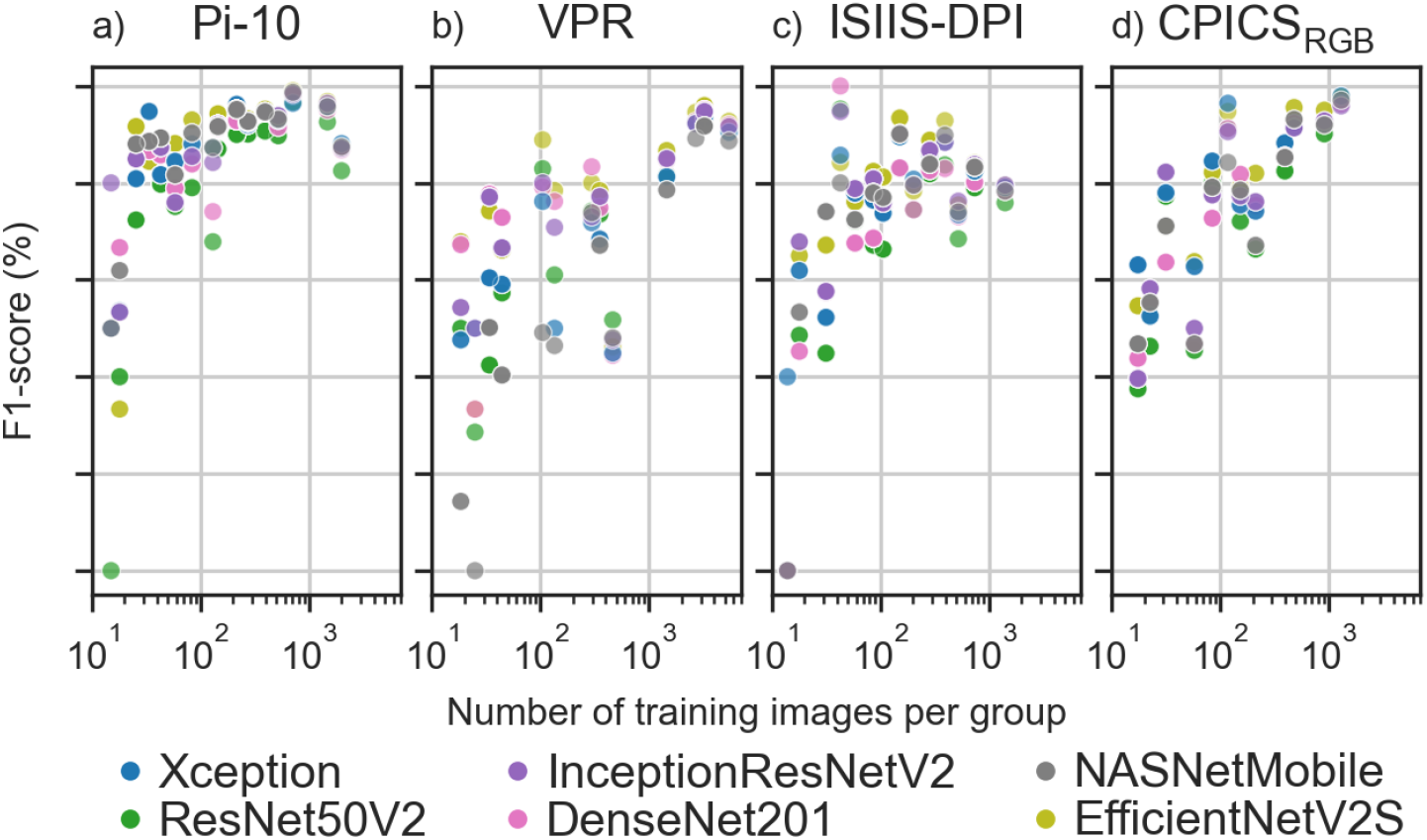
The performance per class for each top-6 model trained on learning set Pi-10 (a), VPR (b), ISIIS-DPI (c) and CPICS_RGB_ (d). The x-axis shows the number of images per class in the training set, and the y-axis shows the F1-score per model per class. Data are binned into logarithmically spaced bins.

Figure 7 shows the F1-score per class of CPICS_RGB_ for the top-6 models. Per model, classes are colour-coded into the 20% best (green) or least (orange) performing quantiles or into neither of these (blue). From the 29 classes in CPICS_RGB_, 13 classes are in the same quantile for all 6 models. This shows that there is little variation in the best and least performing classes between models: classes that perform well in one model tend to perform well in other models. Equivalent figures for the other learning sets are in the appendix (Figure A5, A6 and A7) and show a similar pattern.

**Figure 7:**
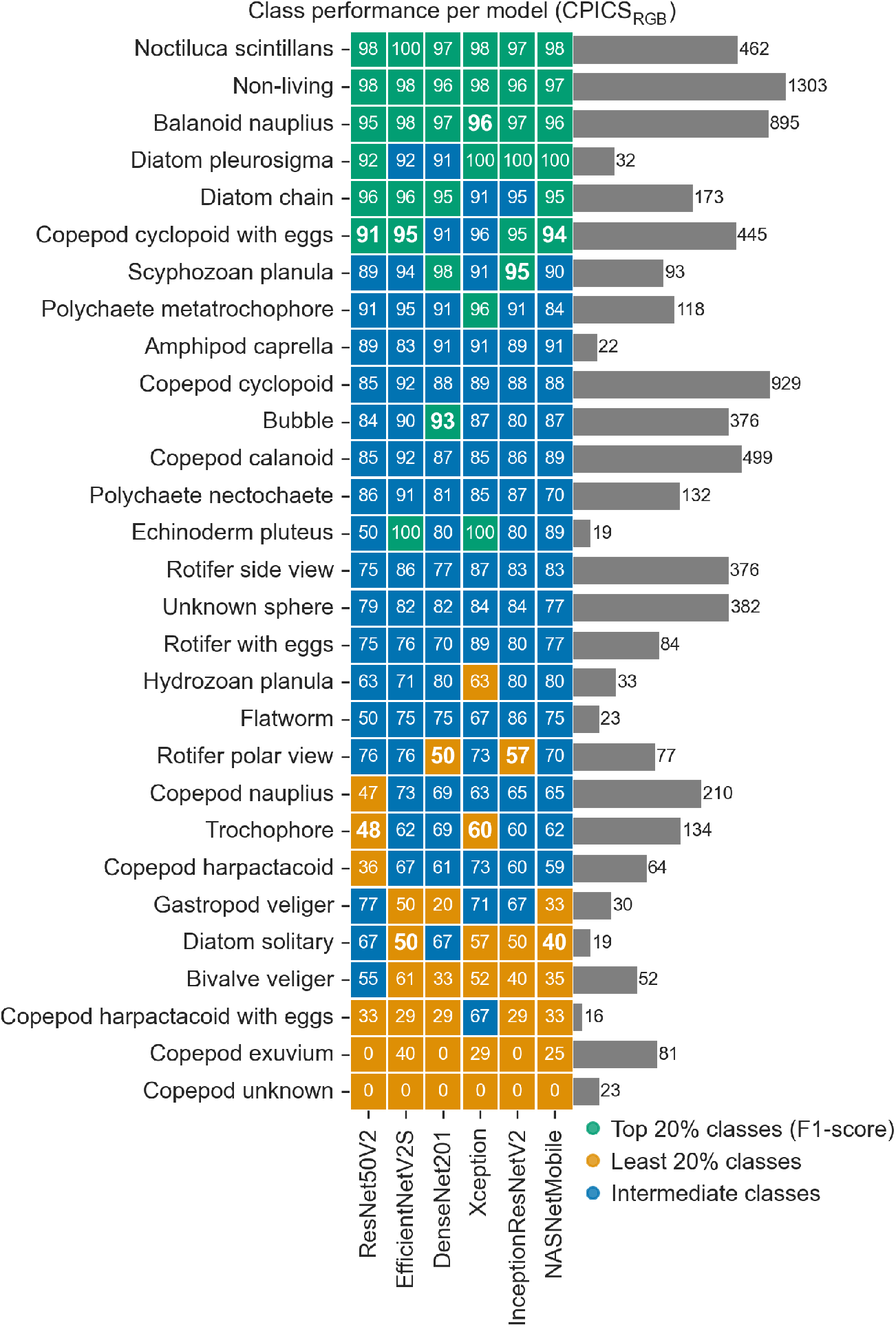
Heatmap of the performance per class for each top-6 model trained on learning set CPICS_RGB_. Colours indicate the quantiles of the F1-score per model: green indicates the 20% best performing classes per model, orange the 20% least performing classes and blue the intermediate classes. Heatmap numbers show the F1-score (as percentage) per class, with the boundary values of the quantiles in boldface. Classes are sorted by the median F1-score of the 6 models. Bars show the number of images per class in the training set. Corresponding figures for the VPR, ISIIS-DPI and Pi-10 are in Figure A5, A6 and A7, respectively.

### Grayscale comparison

For all top-6 models, the performance of the models trained on CPICS_RGB_ is higher than the performance of those trained on the grayscale-transformed equivalent set, CPICS_gray_ (Figure 4, experiment 3). The differences in accuracy and F1-score range from −4.2 to −1.0% and from −1.9 to −7.8%, respectively, with the negative sign indicating that the performance on the RGB set was highest. For the best performing model on CPICS_RGB_, EfficientNetV2S, the performance difference is −1.7% (accuracy) and −2.8% (F1-score). A class-specific comparison shows a negative trend between class size and the grayscale-RGB performance difference (Figure 8): for classes with less than 100 training images, RGB models typically perform up to ∼25% (F1-score) better than their grayscale equivalent, whereas for *>*500 training images, this difference is smaller than 5%. This effect is stronger for DenseNet201, ResNet50V2 and Xception than for EfficientNetV2S, InceptionResNetV2 and NASNetMobile. Overall, we find that colour information increases model performance, especially for classes with few training images.

**Figure 8:**
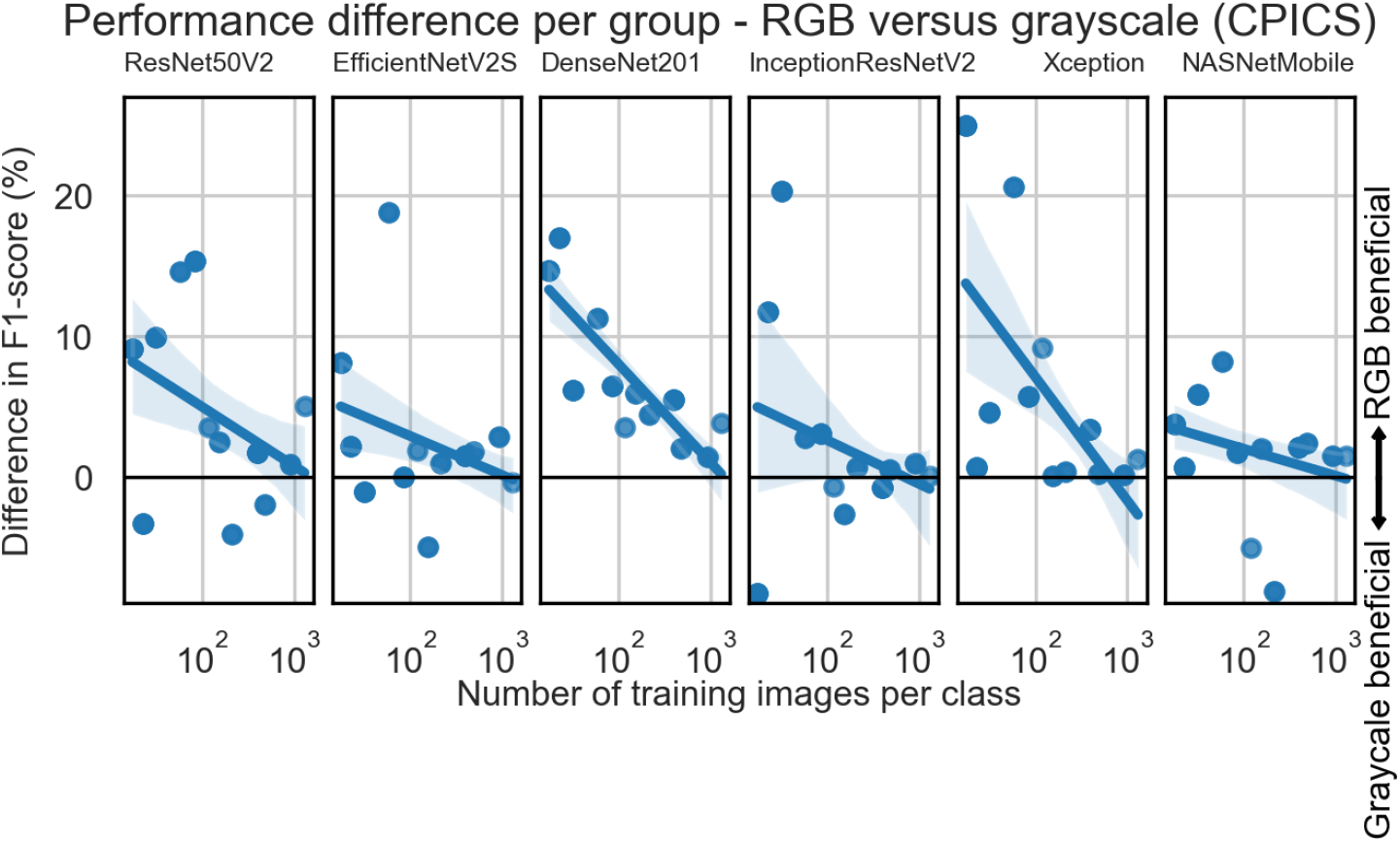
The difference in performance per class between the performance on RGB versus grayscale learning sets from CPICS of the top-6 models. The x-axis shows the number of images per class in the training set, and the y-axis shows the difference in F1-score between the RGB and grayscale performance. Data are binned into logarithmically spaced bins. The lines denote a fitted linear regression model and the shades the 95% confidence interval estimated via bootstrapping.

## Discussion

### Overall performance

The observed differences in network performance on data from CPICS (all models), Pi-10 (top-6 models), VPR (top-6 models) and ISIIS-DPI (top-6 models) show that the choice of CNN architecture can strongly influence the classifier performance. Differences in overall accuracy and F1-score ranged from 4.5 to 7.0%and 7.8 to 19%respectively, with EfficientNetV2S the best performing model in terms of accuracy. Measured in F1-score, EfficientNetV2S is the best (ISIIS-DPI, CPICS_gray_) or second best (VPR, CPICS_RGB_, Pi-10) model. In the latter case, the performance difference with the best performing model was smaller than 1.0%. Our results therefore show that choosing EfficientNetV2S as architecture gives the most reliable model performance.

An additional consideration in the selection of a model, or ensemble of models, is model inference time. Recent studies have focused on ensembling predictions from 4 to 48 CNNs (Lumini and Nanni, 2019; Kyathanahally et al., 2021; Nanni et al., 2023), and such ensembles were shown to achieve higher classification performance than individual models. Another improvement in zooplankton classification performance may be achieved by (ensembles of) Vision Transformers (e.g. Kyathanahally et al., 2022; Maracani et al., 2023; Nanni et al., 2023; Yue et al., 2023; Li et al., 2025). These solutions are computationally heavy. Within single models, we found that the inference time varies 6-fold (Table A4), and naturally, doubling the number of models in a model ensemble leads to a further doubling of the inference time. Substantially increased costs of model inference were also observed for Vision Transformers (Rubbens et al., 2023; Nanni et al., 2025). In use-cases where images need to be classified in real-time, or when tens of millions of images are collected (e.g. Schmid et al., 2023a,b; Pitois et al., 2025; Slocum and Penta, 2025, and Hovenkamp et al., in press), inference time is an important consideration, and using a heavy model (ensemble) may not be feasible.

With the ongoing development of new neural network architectures, new models may emerge that will surpass the performance of the models that we evaluated in this study. Importantly, we found a correlation between the network performance on ImageNet (Deng et al., 2009) and its performance on plankton images (Pearson’s *r* = 0.597 and *p* = 0.0001 for F1-score, Figure 5). This is in agreement with Kornblith et al. (2019), who found a clear correlation between ImageNet accuracy and accuracy on various (non-plankton) data sets in a comparison of 16 CNN architectures. These findings suggest that ImageNet performance is a useful measure when selecting a model architecture for plankton image classification, and it is likely that this also holds for models that have yet to be developed. This implies that for possible new models, an extensive comparison does not need to be done again. Instead, selecting the single best performing, or a handful of the best performing ImageNet models is sufficient in order to identify the best performing model.

### Determinants of classifier performance

We find that performance differences between model architectures are largest for rare classes, varying from ∼20 to 40% F1-score for classes with *<*100 training images, whereas for the most abundant classes (*>*100 training images), all assessed models are within ∼5% F1-score (Figure 6). This shows that selecting a suitable model can substantially increase performance when rare classes are present in the learning set. A different viewpoint is to consider the consistency among models in the classes that perform the best and the least, as Figure 7 shows for CPICS_RGB_.

Achieving acceptable performance on rare classes is a well-known problem in automated image analysis (Brigato et al., 2022) and in plankton image analysis: due to the strong imbalance in the natural abundances of taxonomic classes (Schröder et al., 2019; Eerola et al., 2024; Masoudi et al., 2024), finding images of rare classes in unlabeled data can be a needle-in-a-haystack problem. This is especially profound in (productive) coastal regions with a large abundance of marine snow particles (Forest et al., 2012; Bi et al., 2015; Panäıotis, 2022, this study, Hovenkamp et al., in press) where any living class could be considered rare compared to the number of imaged particles. In addition, the general inverse relation between organism size and abundance (Sheldon et al., 1972) implies that larger organisms such as fish larvae and larger copepods, which may represent pathways to higher trophic levels, are scarce compared to smaller species of zooplankton. As a result, it is common that learning sets are highly unbalanced and that classes of interest may be under-represented. Given the general consensus that more training data per class in general leads to better performance, for these rare classes, classification performance thus tends to lack behind the more abundant classes (Horn and Perona, 2017; Buda et al., 2018), and this can also be seen in Figure 6.

However, one pattern in Figure 7 is that certain classes with few training images (phytoplankton diatom pleurosigma, echinoderm pluteus) are in the best performing quantile for some of the models, whereas some of the most abundant classes (copepod cyclopoid, copepod calanoid, rotifer side view) are in neither of the best performing quantiles, and thus lack behind in performance. Visually, different copepod orders are less distinct than diatoms or echinoderm plutei, and we thus infer that not only the number of training images, but also morphological resemblance plays an important role, which agrees with observations from Kraft et al. (2022) on the automated classification of phytoplankton. Overall, the consistency between models in which classes are in the best and least performing quantile, implies that different model structures do not excel in recognising certain classes compared to other architectures. This indicates that when selecting a model architecture, performance on specific target classes is not a distinguishing factor among architectures.

Second, our results show that performance differences between networks are less pronounced for the Pi-10 than for the other instruments: for the Pi-10, the top-5 models are within a close margin in accuracy (1.5%) and F1-score (2.7%), whereas for the other instruments these differences are at least 2.1% (accuracy) and 6.1% (F1-score). Comparing the image features of the various learning sets visually (Figure 1), we suspect that this is the result of the high image quality of the Pi-10. This is likely a consequence of the different imaging technologies. As mentioned, the Pi-10 is an on-board flow-through system, whereas the other learning sets were collected with in situ instruments (Lombard et al., 2019).

On-board imaging provides several advantages that benefit the image quality. First, despite the similar sampling environment, we generally observed a lower presence of marine snow on Pi-10 images compared to the in situ instruments, which is likely the result of larger particles being broken up by shear stress from the pump. In addition, the on-board set-up results in better control over the object velocity compared to the camera, the object’s distance to the focal plane of the camera, and the amount of light, having less limitations induced by the power supply compared to a stand-alone system. In contrast, with in situ instruments, limited light availability and thus increased exposure time can cause motion blur, and objects may be imaged at a larger distance from the focal plane. Although this is often necessary to obtain a sufficient sampling volume, this compromises the sharpness of some of the images (Lombard et al., 2019). In addition, a larger distance between the camera and the object increases the probability of the object being obscured by marine snow particles. Finally, a notable difference between the Pi-10 and the dark-field imaging systems (CPICS and VPR) that may play a role is the segmentation method. The CPICS and VPR both make use of real-time segmentation methods which discard the non-segmented parts of the images. In our experience, distinguishing features of copepods, such as the urosome or antennules, are commonly discarded during the real-time segmentation procedure of the CPICS and VPR (Figure 1a and 1h) as compared to copepods imaged by the Pi-10 and ISIIS-DPI (Figure 1n and 1s). This benefits automated (and manual) classification of the latter two instruments. We hypothesise then that the high image quality of the Pi-10 resulting from these factors, decreases the need for careful model selection.

In addition, the performance of the Pi-10 on rare classes (30 to 100 images) is substantially higher than for all other instruments that we assessed. We again hypothesise that this is the result of the higher image quality of the Pi-10. This would mean that higher image quality, showing more distinguishable features than images of lower quality, reduces the number of training images that is required to obtain sufficient classification performance. Our findings then imply that there is no general minimum requirement for the number of training images per class, but that this differs per instrument: for the Pi-10, 30 training images per class already result in a F1-score above ∼ 80%, whereas for the CPICS and ISIIS-DPI we estimate this minimum to be ∼100 training images and for VPR, this performance is only consistently reached for the classes with *>*1,000 training images.

An additional benefit of higher image quality is that it could be exploited to increase the taxonomic resolution of the learning set. This is already reflected in the taxonomic resolution of the Pi-10 learning set, which, for example, contains 7 classes of Echinodermata, compared to only 1 for the VPR. This may also affect the classification performance, because distinguishing taxonomically similar classes can be an additional challenge as we discussed earlier this section. How increased taxonomic resolution would affect the sensitivity to model selection and the performance of rare classes for the Pi-10, would be an interesting topic for future research.

### The value of colour information

Colour information increased the performance of the best performing classifier on CPICS_RGB_ (EfficientNetV2S) with 1.7% (accuracy) and 2.8% (F1-score). The importance of colour information for CNN classification has gained, to our knowledge, surprisingly little attention. For non-zooplankton data sets, CNNs are believed to rely most on shape (e.g. Kubilius et al., 2016) and texture (e.g. Geirhos et al., 2019), and the dependency on colour information may differ per data set (Singh et al., 2020). A previous study on zooplankton images has shown that converting images to grayscale before running an RGB-trained CNN classifier decreases the accuracy by 4.5% as compared the RGB equivalent of the same set (Li et al., 2022). This shows that a network that is trained on colour images indeed makes use of the colour information. This is in line with our findings here, but we extend this conclusion by comparing classifiers trained on CPICS colour images to classifiers *trained* on grayscale-converted images. The observed performance increase is strongest (∼25% F1-score) for classes with low (*<*100) numbers of training images, but the increase in performance decreases to *<*5% for the classes with *>*500 images. As a result, the overall performance increase (2.8% F1-score) is smaller than expected, but when interested in finding rare classes, colour information provides a clear benefit.

This result poses the question whether recording colour information with in situ zooplankton imaging instruments is worth the increased data handling. For a stand-alone imaging instrument (i.e. without a data connection to the deck), the amount of data that can be recorded during a deployment is limited. Storing colour images (three channels) instead of grayscale images (single channel) increases the data stream and storage, and therefore compromises other factors that increase the amount of recorded data, such as the image resolution or the volume of seawater that is imaged during a deployment. Consequently, as was pointed out by Lombard et al. (2019), a lower sampling volume decreases the upper limit on the size of imaged zooplankton taxa. Of course, colour information of zooplankton images can provide other advantages than automated taxonomic identification alone, as it can also be used to recognise additional features (Campbell et al., 2020). In addition, for the case of phytoplankton classification, colour (as recorded by e.g. FlowCAM, Sieracki et al., 1998) might prove to be a more valuable feature than for the zooplankton classes assessed in this study. This would be an interesting topic for future research. When deploying or developing new zooplankton imaging instruments, the value of colour is an important consideration to take into account: with an overall performance increase of only 1.7 to 2.8% (accuracy resp. F1-score) on our best performing classifier when going from grayscale to RGB images, increasing the frame rate and thus sampling volume appears more beneficial than recording colour.

An interesting consequence of this finding may be the use of domain adaptation techniques (as suggested by e.g. Eerola et al., 2024; Batrakhanov et al., 2024; Ciranni et al., 2024) to construct classifiers that can benefit from training data from multiple instruments, which would be an interesting direction for future research. With domain adaptation techniques, the growing number of publicly available annotated data sets (e.g. EcoTaxa, Picheral et al., 2017), which are mostly in grayscale, could theoretically aid in classifying (newly collected) colour images, for which the amount of labeled data is limited. However, combining colour and grayscale instruments in a single classifier is not trivial: grayscale training images cannot learn a classifier to utilise colour information, but in case the colour information would not be used, potentially useful information is thus discarded. The trade-off that our results suggest between number of training images and the advantage of colour information, implies that discarding colour information when it leads to substantially more training images may nevertheless increase a classifier’s performance on rare classes.

## Conclusion

Our study shows that for all data sets that we tested, EfficientNetV2S is a reliable choice to achieve good model performance. Moreover, we found a correlation between ImageNet performance and performance on zooplankton images, which extends our findings to network architectures that were not assessed here: a model that performs well on ImageNet is expected to perform well for zooplankton classification.

In addition, we investigated various factors that affect the performance of CNNs for zooplankton classification. We found that the high image quality of the Pi-10 decreases the need for careful model selection, and that colour information increases the performance of the best performing classifier on CPICS_RGB_ (EfficientNetV2S) with 1.7 to 2.8% (accuracy resp. F1-score), which is less than expected and questions the advantage of recording colour information for zooplankton.

Finally, we provide insights to improve the classification performance of rare classes. The presence of rare classes is common in many zooplankton studies: in turbid conditions in shelf seas, which are often highly productive systems, marine snow can be up to 99% of the data. This makes any living class in the learning set rare, because manually labeling these is a needle-in-the-haystack problem. When such rare classes are present, we found that colour information and the high image quality of the Pi-10 reduces the number of training images that are needed to achieve good performance. For the Pi-10, our results indicate that an F1-score above 80% can be reached consistently already with 30 training images per class, but for the CPICS and ISIIS-DPI this performance required roughly ∼100 training images, and for the VPR *>*1,000. Moreover, performance differences between model architectures were largest for the less abundant classes, which implies that when these are present, careful model selection is most beneficial.

## Data Availability Statement

The annotated VPR set was published online (Flanders Marine Institute (VLIZ) Belgium, 2023, https://doi.org/10.14284/607), and the annotated ISIIS-DPI set is available at https://doi.org/10.5281/zenodo.20762202. The CPICS and Pi-10 sets and all Python code are available at https://doi.org/10.5281/zenodo.21074144.

## Acknowledgments

This study was made possible through collaborative funding between Utrecht University, the Netherlands, and the Royal Netherlands Institute for Sea Research (NIOZ). Pi-10 data was obtained within a project funded by the European Maritime and Fisheries and Aquaculture Fund (EMFAF). This study was supported by data and infrastructure provided by VLIZ as part of the Flemish contribution to LifeWatch.

## Author Contribution Statement

PH, AFvdS and DvO conceived and designed the study. LvW, DvO, AO and PH collected the field data. LvW, AO and PH manually labelled the training images. PH carried out the numerical experiments. AFvdS, DvO, LvW, AO and PH interpreted the results. PH wrote the manuscript with the support of AFvdS and DvO and with significant input from all authors. AFvdS and DvO supervised the study and contributed equally to this work.

## Appendix

**Figure A1:**
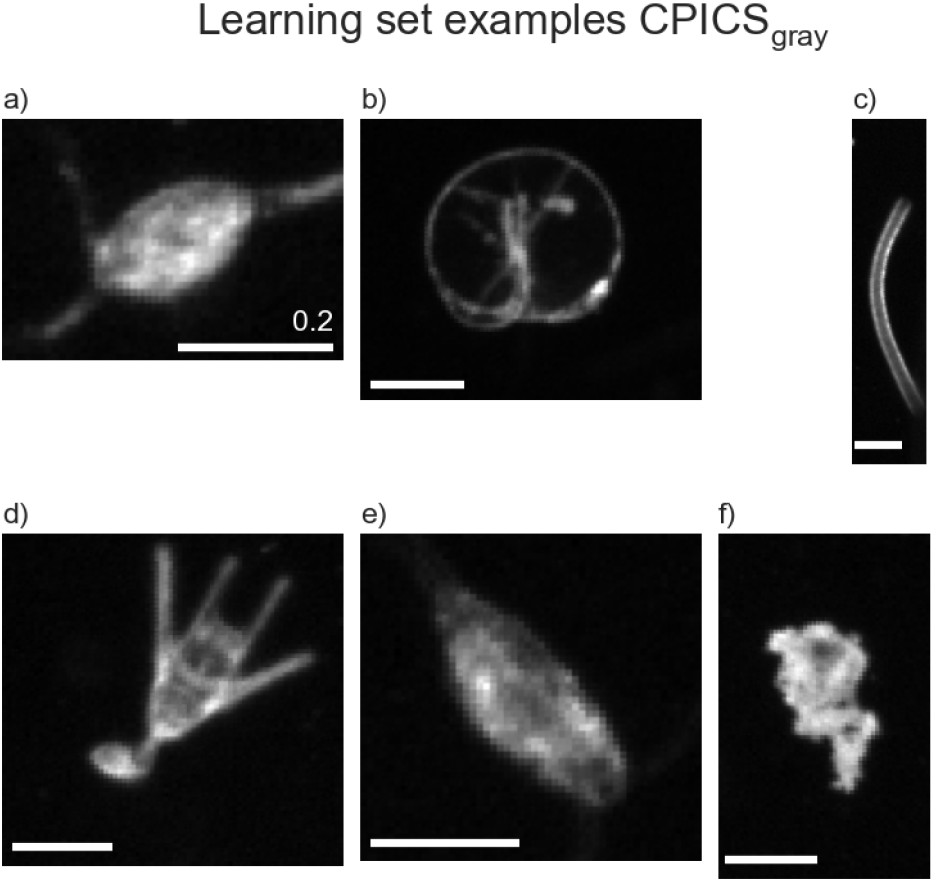
Example of grayscale-converted images from CPICS_RGB_, which form the learning set CPICS_gray_. Species names and original RGB-version of the ROIs are in Fig. 1. Scale bars denote a size of 2 mm.

**Figure A5:**
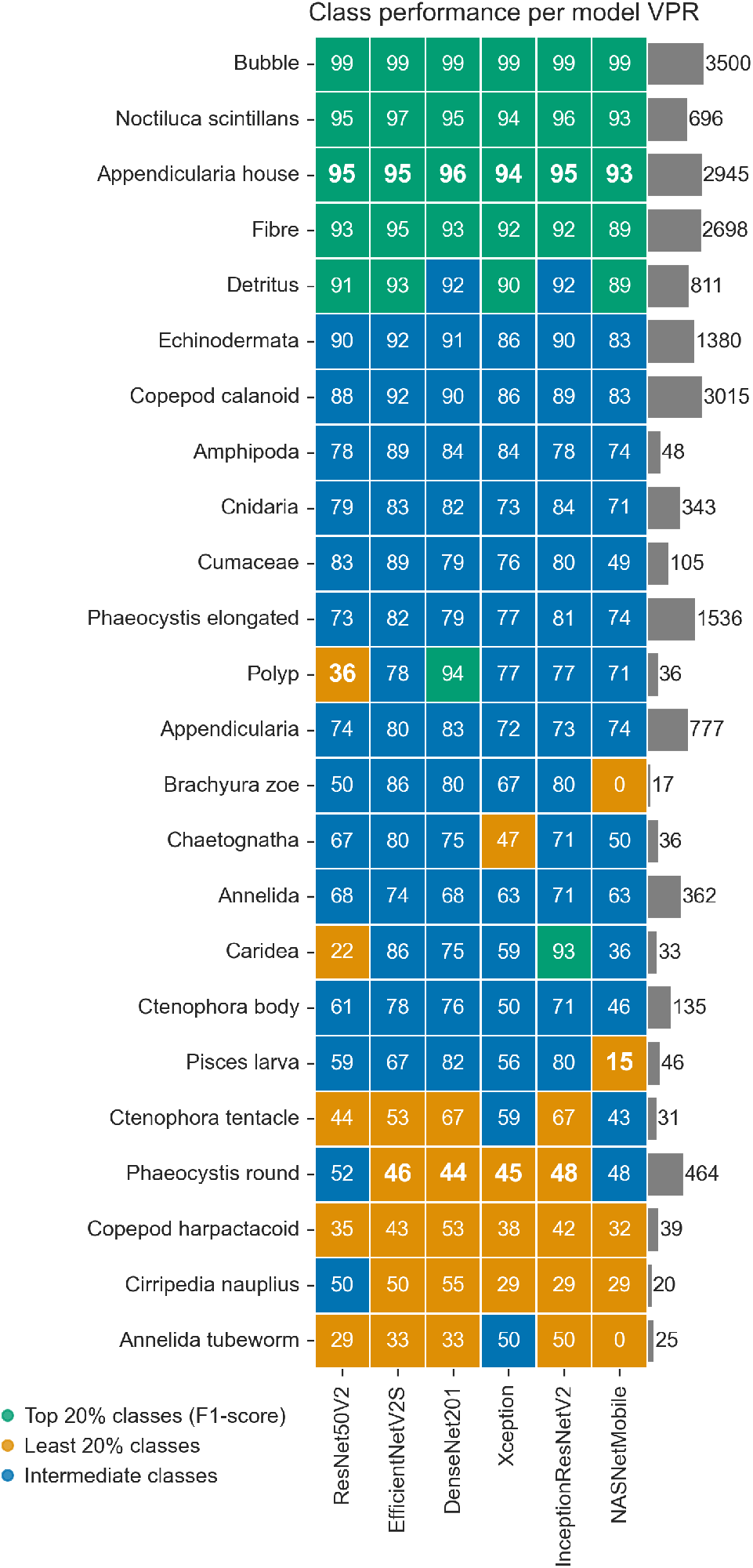
Heatmap of the performance per class for each top-6 model trained on learning set of VPR. Colours indicate the quantiles of the F1-score per model: green indicates the 20% best performing classes per model, orange the 20% least performing classes and blue the intermediate classes. Heatmap numbers show the F1-score (as percentage) per class, with the boundary values of the quantiles in boldface. Classes are sorted by the median F1-score of the 6 models. Bars show the number of images per class in the training set.

**Figure A6:**
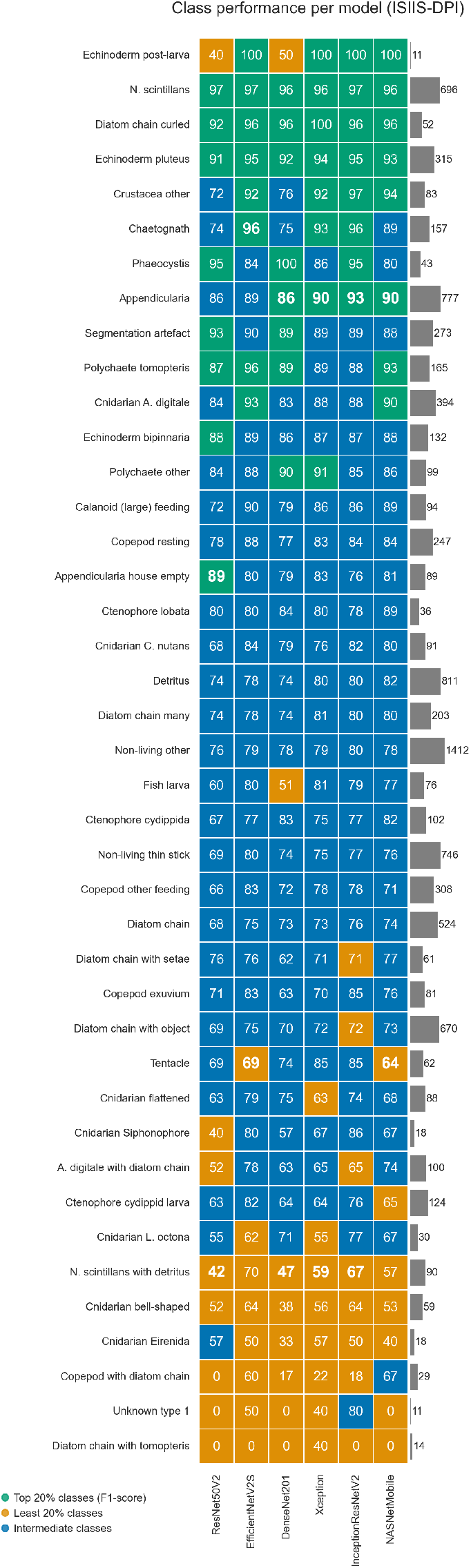
Heatmap of the performance per class for each top-6 model trained on learning set of ISIIS. The rest of the description is similar to Fig. A5.

**Figure A7:**
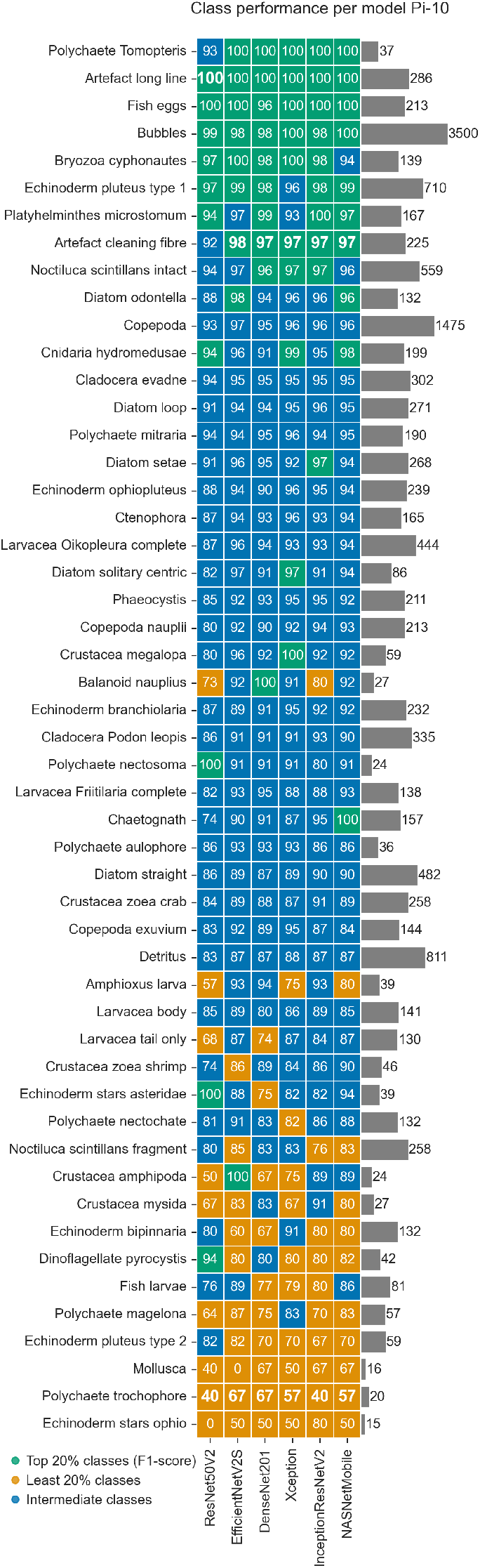
Heatmap of the performance per class for each top-6 model trained on learning set of Pi-10. The rest of the description is similar to Fig. A5.

---

**Table A2:** Properties of the Convolutional Neural Network (CNN) architectures that were used in this study.

| *Architecture* | *Image size (pixels)* | *Trainable parameters (million)* | *ImageNet top-1 accuracy (%)* | *Reference* |
| --- | --- | --- | --- | --- |
| DenseNet121 | 224 | 7.0 | 75.0 | Huang et al. (2017) |
| DenseNet169 | 224 | 12.5 | 76.2 |  |
| DenseNet201 | 224 | 18.1 | 77.3 |  |
| EfficientNetV2B0 | 224 | 5.9 | 78.7 | Tan and Le (2021) |
| EfficientNetV2B1 | 240 | 6.9 | 79.8 |  |
| EfficientNetV2B2 | 260 | 8.7 | 80.5 |  |
| EfficientNetV2B3 | 300 | 12.9 | 82.0 |  |
| EfficientNetV2S | 384 | 20.2 | 83.9 |  |
| InceptionResNetV2 | 299 | 54.3 | 80.3 | Szegedy et al. (2017) |
| InceptionV3 | 299 | 21.8 | 77.9 | Szegedy et al. (2016) |
| MobileNetV2 | 224 | 2.3 | 71.3 | Sandler et al. (2018) |
| NASNetMobile | 224 | 4.3 | 74.4 | Zoph et al. (2018) |
| ResNet50V2 | 224 | 23.6 | 76.0 | He et al. (2016) |
| ResNet101V2 | 224 | 42.6 | 77.2 |  |
| ResNet152V2 | 224 | 58.2 | 78.0 |  |
| VGG16 | 224 | 134.4 | 71.3 | Simonyan and Zisserman (2015) |
| VGG19 | 224 | 139.7 | 71.3 |  |
| Xception | 299 | 20.9 | 79.0 | Chollet (2017) |

**Table A3:**
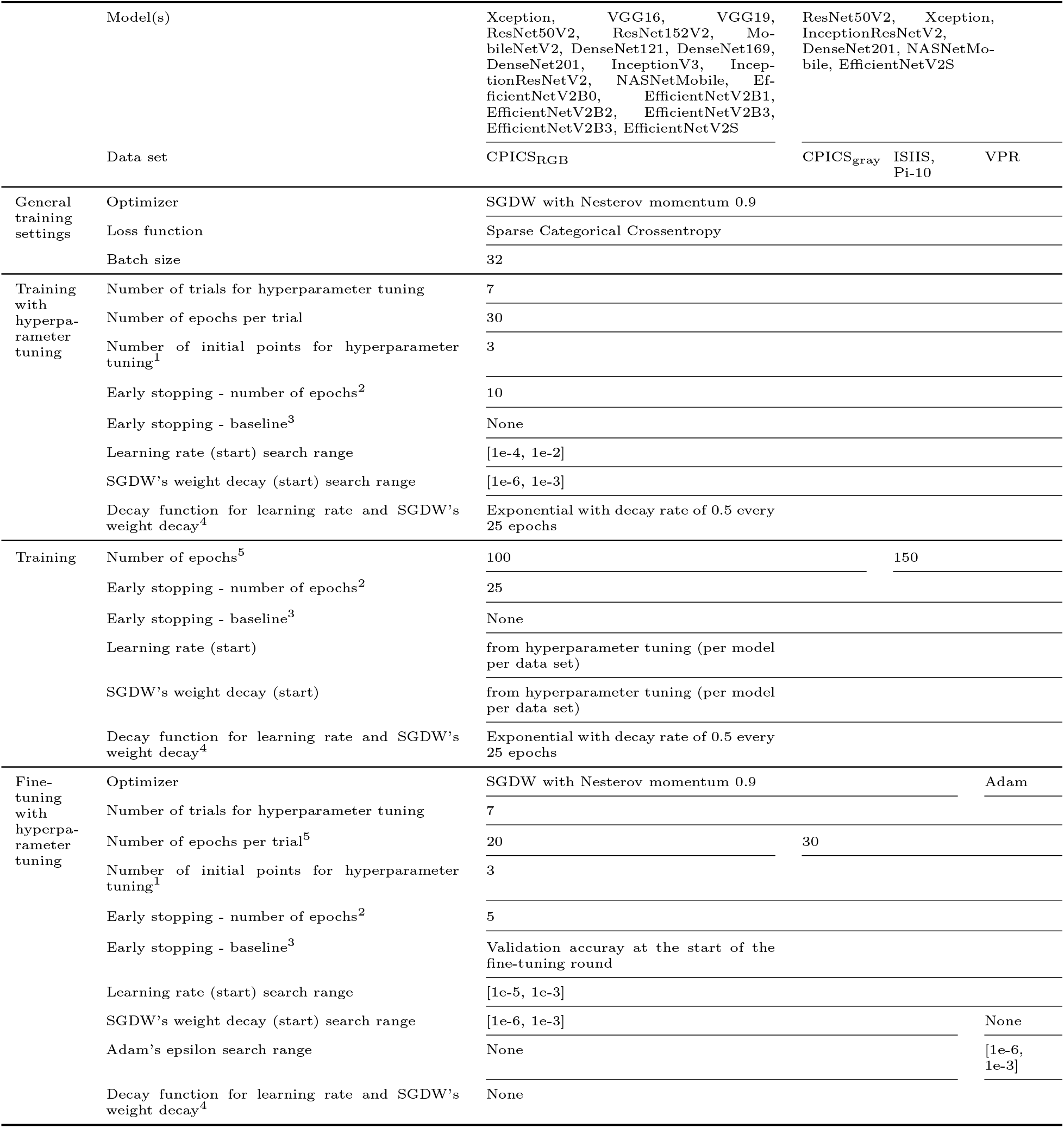
The training parameters that were used for training the CNN classifiers per data set per model. ^1^the number of trials for which the assessed hyperparameter values are randomly drawn from the search space. After these initial points, the next hyperparameters values are determined using Bayesian optimization (caption continues on next page). ^2^a trial is aborted if the validation accuracy has not improved for this number of epochs. ^3^a trial is aborted if the validation accuracy is lower than this baseline after the number of epochs of’Early stopping - number of epochs’. ^4^if specified, an exponential decay function determines the learning rate and SGDW’s weight decay (if applicable) for every epoch. When applicable, the specified values for learning rate and weight decay are the values in the first epoch. ^5^for all results it was checked if the training process had converged based on the training history. If this was not the case, the number of epochs was increased.

**Table A4:**
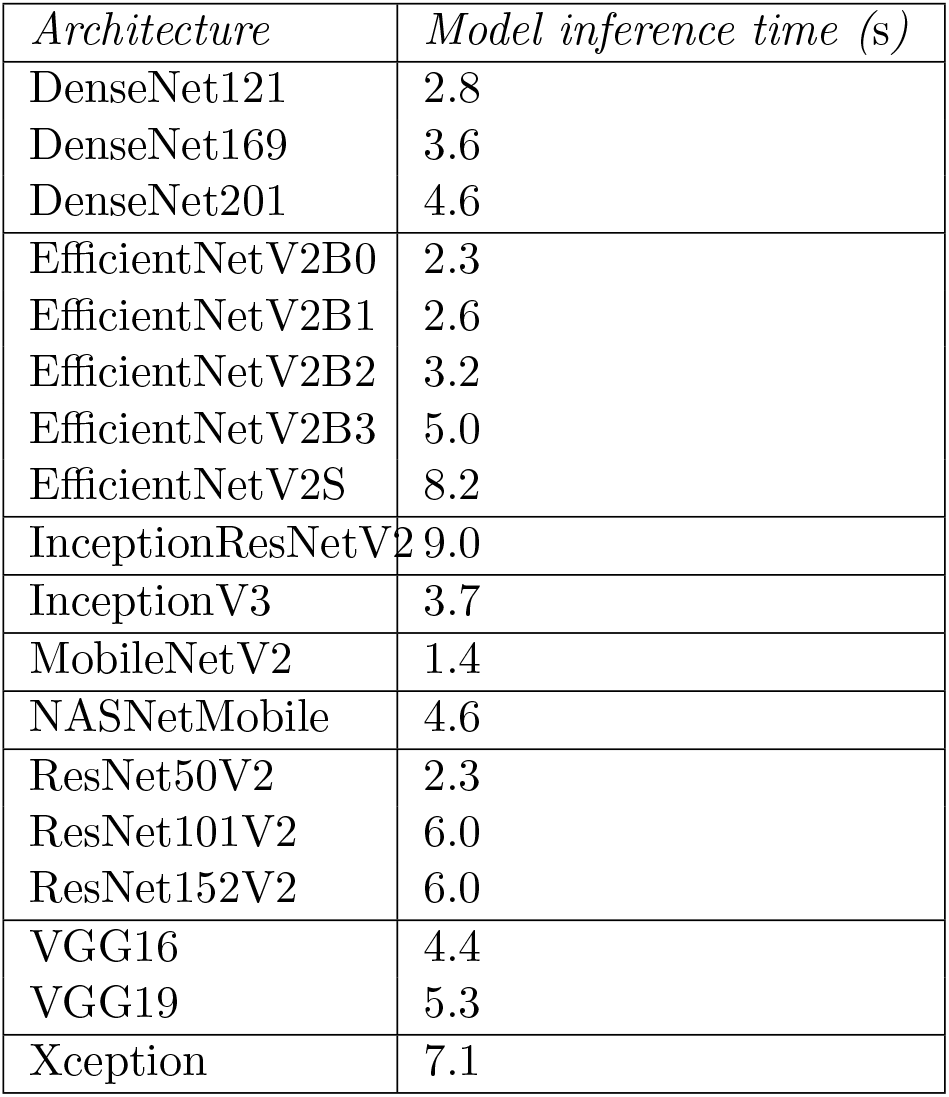
Model inference time of the Convolutional Neural Network (CNN) architectures that were used in this study, as evaluated on 1,780 images of CPICS_RGB_ averaged over 20 trials on a MacStudio (M1 Ultra chip with 64GB memory).

| *Architecture* | *Model inference time (s)* |
| --- | --- |
| DenseNet121 | 2.8 |
| DenseNet169 | 3.6 |
| DenseNet201 | 4.6 |
| EfficientNetV2B0 | 2.3 |
| EfficientNetV2B1 | 2.6 |
| EfficientNetV2B2 | 3.2 |
| EfficientNetV2B3 | 5.0 |
| EfficientNetV2S | 8.2 |
| InceptionResNetV2 | 9.0 |
| InceptionV3 | 3.7 |
| MobileNetV2 | 1.4 |
| NASNetMobile | 4.6 |
| ResNet50V2 | 2.3 |
| ResNet101V2 | 6.0 |
| ResNet152V2 | 6.0 |
| VGG16 | 4.4 |
| VGG19 | 5.3 |
| Xception | 7.1 |

